# Divergent behavioral consequences of manipulations enhancing pyramidal neuron excitability in the prelimbic cortex

**DOI:** 10.1101/2020.06.04.134486

**Authors:** Timothy R. Rose, Ezequiel Marron Fernandez de Velasco, Baovi N. Vo, Megan E. Tipps, Kevin Wickman

**Author notes:** equal contributors (Timothy R. Rose, Ezequiel Marron Fernandez de Velasco). **Corresponding author**, Kevin Wickman, PhD, Department of Pharmacology, University of Minnesota, 6-120 Jackson Hall, 312 Church Street SE, Minneapolis, MN 55455.

## Abstract

**Background:** Drug-induced neuroadaptations in the prefrontal cortex are thought to underlie impaired executive functions that reinforce addictive behaviors. Repeated cocaine exposure increased layer 5/6 pyramidal neuron excitability in the mouse prelimbic cortex (PL), an adaptation attributable to a suppression of G protein-gated inwardly rectifying K^+^ (GIRK/Kir3) channel activity. GIRK channel suppression in the PL of drug-naïve mice enhanced the motor-stimulatory effect of cocaine. The impact of cocaine on PL GABA neurons, key pyramidal neuron regulators, and the behavioral relevance of increased PL pyramidal neuron excitability, remain unclear.

**Methods:** The effect of repeated cocaine on mouse layer 5/6 PL GABA neurons was assessed using slice electrophysiology. Adaptations enhancing PL pyramidal neuron excitability were modeled in drug-naïve mice using persistent viral Cre ablation and acute chemogenetic approaches. The impact of these manipulations on PL-dependent behavior was assessed in motor activity and trace fear conditioning tests.

**Results:** Repeated cocaine treatment did not impact GIRK channel activity in, or excitability of, layer 5/6 PL GABA neurons. GIRK channel ablation in PL pyramidal neurons enhanced the motor-stimulatory effect of cocaine but did not impact baseline activity or fear learning. In contrast, direct or indirect chemogenetic activation of PL pyramidal neurons increased baseline and cocaine-induced motor activity and disrupted fear learning. These effects were mirrored by chemogenetic activation of PL pyramidal neurons projecting to the ventral tegmental area.

**Conclusions:** Manipulations enhancing the excitability of PL pyramidal neurons, including those projecting to the VTA, recapitulate behavioral hallmarks of repeated cocaine exposure.

## INTRODUCTION

The medial prefrontal cortex (mPFC) plays a crucial role in cognition and regulation of motivated behavior (1, 2). The mPFC provides glutamatergic input to several brain regions including the ventral tegmental area (VTA), basolateral amygdala (BLA), and nucleus accumbens (NAc) (3, 4), and these projections have been linked to key facets of cocaine addiction (1, 2, 5). For example, cocaine exposure increases glutamate release in the NAc and VTA (6), and these increases and associated cocaine-induced neuroadaptations and drug-seeking behavior can be blocked by mPFC inactivation (7–11). In addition, cocaine-induced adaptations in mPFC projections are critical for the development and expression of locomotor sensitization, a phenomenon sharing anatomic and neurochemical features with craving (5).

The mPFC consists of subregions including the cingulate, prelimbic (PL), infralimbic (IL), and orbitofrontal cortices (1, 2). Numerous studies have highlighted the role of the PL in regulating addiction-related behaviors and cognition (2, 12–16). For example, PL lesions prevent the induction and expression of cocaine-induced locomotor sensitization, and PL inactivation decreases the reinstatement of cocaine-seeking behavior (17–19). PL activity is also necessary for associative learning (20, 21), which is dysregulated following repeated cocaine exposure (22). In rodent trace fear conditioning studies, for example, persistent firing in the PL during the trace interval, the period separating the auditory cue and delivery of a footshock, is critical for acquisition of fear learning (23–32). Indeed, trace fear learning is prevented by optogenetic silencing of the PL during the trace interval (32).

The PL contains excitatory pyramidal neurons (~85%) and GABAergic interneurons (~15%) (33, 34). Pyramidal neurons, particularly those from layers 5 and 6, are the primary projection neurons, while GABA neurons regulate pyramidal neuron excitability (3, 4). Repeated cocaine exposure triggers an array of adaptations that increase PL pyramidal neuron excitability (35–42). Repeated cocaine also reduces GABAergic neurotransmission in PL pyramidal neurons via suppression of presynaptic GABA release (43), and blunting of postsynaptic GABA_A_R- and GABA_B_R-mediated signaling (40, 43, 44). At present, the behavioral relevance of elevated PL pyramidal neuron excitability is not well-understood.

Previously, we reported that a cocaine sensitization regimen increased layer 5/6 PL pyramidal neuron excitability in mice, and that this adaptation correlated with reduced G protein-gated inwardly rectifying K^+^ (GIRK/Kir3) channel activity (40). Viral suppression of GIRK channel activity in the PL of drug-naïve mice increased the motor-stimulatory effect of cocaine. The potential impact of repeated cocaine on PL GABA neuron excitability is unclear, however, as are the behavioral consequences of elevated PL pyramidal neuron excitability. Here, we examined the impact of repeated cocaine on layer 5/6 PL GABA neurons, and used neuron-specific viral approaches to probe the behavioral impact of distinct manipulations that persistently or acutely enhance PL pyramidal neuron excitability.

## MATERIALS AND METHODS

### Animals

All experiments were approved by the University of Minnesota Institutional Animal Care and Use Committee. The generation of *Girk1^-/-^*, *Girk2^-/-^*, and *Girk1^fl/fl^* mice was described previously (45–47). GAD67GFP mice were provided by Dr. Takeshi Kaneko (48). CaMKIICre (B6.Cg-Tg(Camk2a-cre)T29-1Stl/J) and GADCre (B6N.Cg-*Gad2^tm2(cre)Zjh^*/J) lines were purchased from The Jackson Laboratory (Bar Harbor, ME) and were maintained by backcrossing against the C57BL/6J strain. Cre(+) and/or Cre(-) offspring were used in some experiments. Male C57BL/6J mice were purchased for some studies. Mice were maintained on a 14:10 h light/dark cycle and were provided *ad libitum* access to food and water.

### Chemicals

Baclofen, barium chloride, picrotoxin, and kynurenic acid were purchased from Sigma (St. Louis, MO). CGP54626, clozapine-N-oxide (CNO), and tetrodotoxin were purchased from Tocris (Bristol, UK). Cocaine was obtained through Boynton Health Pharmacy at the University of Minnesota.

### Viral vectors

pAAV-hSyn-DIO-hM3Dq(mCherry) (Addgene plasmid #44361), pAAV-hSyn-DIO-hM4Di(mCherry) (Addgene plasmid #44362), and pAAV-hSyn-DIO-mCherry (Addgene plasmid #50459) were gifts from Dr. Bryan Roth. pAAV-CaMKIIα-hM3Dq(mCherry) and pAAV-CaMKIIa-mCherry plasmids were generated by the University of Minnesota Viral Vector and Cloning Core (VVCC; Minneapolis, MN) using standard cloning techniques and pAAV-CaMKIIα-hChR2(C128S/D156A)-mCherry (Addgene plasmid #35502, a gift from Dr. Karl Deisseroth) as the backbone. Similarly, pAAV-mDlx-hM4Di(mCherry), pAAV-mDlx-mCherry and pAAV-mDlx-tdTomato were generated using pAAV-mDlx-GCaMP6f-Fishell-2 (Addgene plasmid #83899, a gift from Dr. Gordon Fishell) as the source of the mDlx promoter/enhancer. pAAV-hSyn-Cre-GFP (Addgene plasmid #68544, a gift from Dr. Eric Nestler) was packaged into AAV2retro. AAV8-CaMKIIa-Cre(mCherry) was purchased from the University of North Carolina Vector Core (Chapel Hill, NC). All other viral vectors were packaged in AAV8 serotype by the University of Minnesota VVCC (Minneapolis, MN); all viral titers were between 3.5 x 10^12^ - 2.2 x 10^14^ genocopies/mL.

### Intracranial viral manipulations

Intracranial infusion of virus (400 nL per side) in mice (7-8 wk) was performed as described (49), using the following coordinates (in mm from bregma: AP, ML, DV): PL (+2.50, ±0.45, −1.60), BLA (−1.50, ±3.35, −4.70), NAc (+1.50, ±1.00, −4.50), and VTA (−2.60, ±0.65, - 4.70). After surgery, animals were allowed 3-4 wk (chemogenetic studies) or 4-5 wk (Cre ablation or retrograde chemogenetic studies) for full recovery and viral expression before electrophysiological or behavioral assessments. The scope and accuracy of targeting was assessed using fluorescence microscopy. Brightfield and fluorescent images were overlaid and evaluated using the *Mouse Brain Atlas* (50). Targeting coordinates and viral loads yielded extensive coverage of the PL along the rostro-caudal axis, with limited spread into the anterior cingulate (cg), medial orbital, or IL cortices. Only data from mice in which >70% of viral-driven bilateral fluorescence was confined to the PL were included in the final analysis. To evaluate the targeting fidelity of AAV8/CaMKIIα- and AAV8/mDlx-based vectors, AAV8-CaMKIIα-mCherry or AAV8-mDlx-mCherry vectors were infused into the PL of GAD67GFP(+) mice. After a 2-wk period, brains were fixed with 4% paraformaldehyde, coronal sections (50 micron) were obtained by sliding microtome, and images of viral-driven mCherry and GFP fluorescence were acquired. Quantification of cells expressing mCherry, GFP, or both (overlap) was performed with ImageJ software (51).

### Slice electrophysiology

Baclofen-induced somatodendritic currents were recorded in layer 5/6 PL neurons, as described (52). For rheobase assessments, cells were held at 0 pA in current-clamp mode and given 1-s current pulses, beginning at −60 pA and increasing in 20 pA increments. Rheobase was identified as the injection step at which initial spiking was elicited. For PL GABA neuron recordings, rheobase was measured prior to and after perfusion of baclofen (200 μM). For chemogenetic experiments, resting membrane potential and rheobase were assessed prior to and after bath perfusion of CNO (10 μM). Spontaneous inhibitory postsynaptic currents (sIPSCs) were recorded and analyzed, as described (49).

### Behavioral testing

Adult mice (10-13 wk) were evaluated in open-field motor activity and trace fear conditioning tests. For motor activity studies, mice were acclimated to handling, injection, and open field chambers for 2-4 d prior to testing. For GIRK ablation experiments, distance traveled during the 60-min interval after saline injection on the final acclimation day was taken as baseline activity. Distance traveled after injection of cocaine (15 mg/kg IP) the next day was taken as cocaine-induced activity. For chemogenetic studies, CNO (2 mg/kg IP) was administered 30 min prior to saline injection and placement in the open field; distance traveled over the next 60-min was taken as baseline activity. Subsequently (2-4 d later), subjects were injected with CNO (2 mg/kg IP) 30-min prior to cocaine (15 mg/kg IP); distance traveled over the next 60-min interval was taken as cocaine-induced activity. In studies involving AAV8/mDlx-based vectors, separate cohorts of mice underwent baseline or cocaine-induced activity testing.

For trace fear conditioning studies, mice were acclimated to handling and testing room for 1-2 d prior to testing. The 6.5-min conditioning session (Day 1) involved 2 pairings of a 30-s auditory cue (65 dB white noise) and a 2-s footshock (0.5 mA), separated by a 30-s trace interval. For chemogenetic studies, CNO (2 mg/kg IP) was only administered once, 30 min prior to conditioning on Day 1. Cue recall was assessed on Day 3, with chambers reconfigured using a white plastic insert to cover the bar floor and a black tent insert to alter the size, shape, and color of the environment. Inserts were also cleaned with 0.1% acetic acid instead of ethanol to provide a distinct olfactory cue. Freezing was monitored throughout the 15-min test period, divided into 5 x 3-min bins that included 2 x 3-min auditory cue presentations. For projection-specific manipulations, motor activity testing was performed 3-12 d after the trace fear conditioning study.

### Data analysis

Data are presented as the mean ± SEM. Statistical analyses were performed using Prism 8 software (GraphPad Software, Inc.; La Jolla, CA). Unless specifically noted, all studies involved balanced groups of male and females. While sex was included as a variable in preliminary analyses, no impact of sex was observed on any measure and data from male and females were pooled. Pooled data were analyzed by paired and unpaired student’s *t* test, Mann-Whitney test, one-way ANOVA, twoway ANOVA, two-way repeated-measures ANOVA, and mixed-effects model (REML), as appropriate. Pairwise comparisons were performed using Bonferroni’s *post hoc* test, if justified. Data points that fell outside of the group mean by more than 2 standard deviations were excluded from analysis; this resulted in the exclusion of 11 data points across the entire study. Differences were considered significant when *P*<0.05.

## RESULTS

### Impact of repeated cocaine on PL GABA neurons

Repeated cocaine administration in mice suppressed GIRK channel activity in layer 5/6 PL pyramidal neurons, and viral suppression of GIRK channel activity in the mouse PL increased cocaine-induced motor activity (40). This manipulation was not selective for pyramidal neurons, however, and GABAergic neurons in the mPFC regulate pyramidal neuron excitability (3, 4). Since psychostimulant exposure suppressed GIRK-dependent signaling in VTA GABA neurons (53), we tested whether layer 5/6 PL GABA neurons express GIRK channels, and if so, whether repeated cocaine exposure alters GIRK-dependent signaling in, or excitability of, these neurons.

We used GAD67GFP(+) transgenic mice (48) to facilitate targeted electrophysiological analysis of layer 5/6 PL GABA neurons (**Fig. 1A**). The GABA_B_R agonist baclofen evoked an outward current in PL GABA (GFP-positive) neurons that correlated with decreased input resistance; no sex difference was detected (*t*_17_=0.73, *P*=0.47). Baclofen-induced responses were suppressed by 0.3 mM external Ba^2+^, consistent with GIRK channel activation (**Fig. 1B,C**). Indeed, PL GABA neurons lacking GIRK1 or GIRK2 exhibited diminished baclofen-induced currents (**Fig. 1D,E**). GIRK1 or GIRK2 abation did not impact rheobase (**Fig. 1F**), but did blunt the baclofen-induced increase in rheobase (**Fig. 1G**). Thus, layer 5/6 PL GABA neurons express a GIRK channel, formed by GIRK1 and GIRK2, that mediates most of the postsynaptic GABABR-dependent influence on excitability. We next subjected GAD67GFP(+) mice to a locomotor sensitization regimen involving daily injections of cocaine (15 mg/kg IP) or saline over 5 consecutive days, as described (40). Subsequently (1-2 d later), we measured resting membrane potential (RMP), rheobase, and baclofen-induced currents in layer 5/6 PL GABA neurons. Repeated cocaine had no impact on baclofen-induced current amplitude (**Fig. 1H,I**), RMP (**Fig. 1J**), or rheobase (**Fig. 1K**).

**Figure 1.**
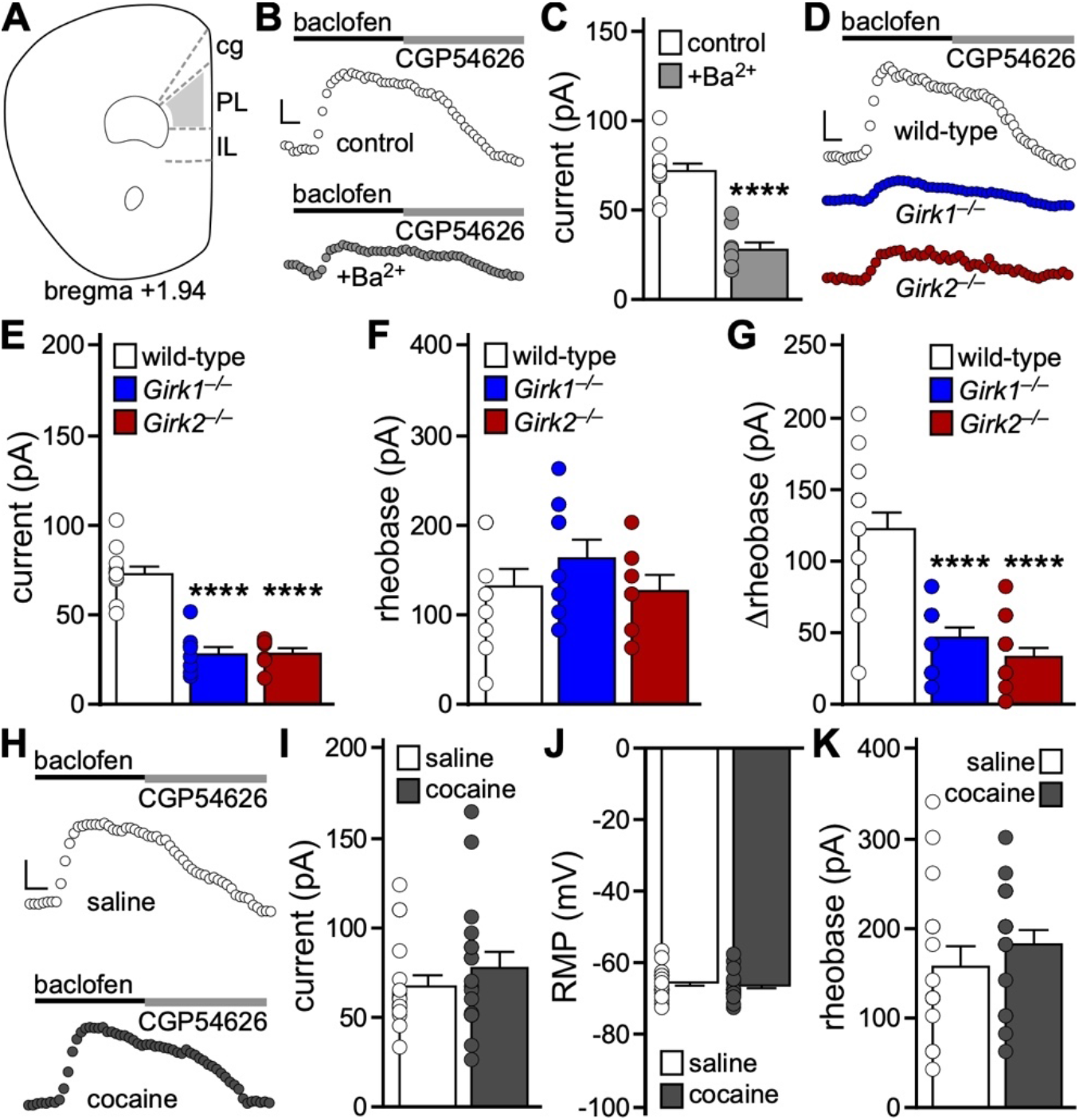
Impact of repeated cocaine exposure on layer 5/6 PL GABA neurons. **A.** Schematic highlighting the PL, and adjacent cingulate (cg) and infralimbic (IL) cortices. GFP-positive (GABA) neurons in layer 5/6 of the PL, in slices from GAD67GFP(+) mice, were targeted for analysis. **B.** Somatodendritic currents (V_hold_=-60 mV) evoked by baclofen (200 μM) in GABA neurons from GAD67GFP(+) mice, in the absence and presence of external 0.3 mM Ba^2+^. Currents were reversed by the GABA_B_R antagonist CGP54626 (2 μM). Scale: 25 pA/50 s. **C.** Baclofen-induced currents in GABA neurons from GAD67GFP(+) mice, in the absence and presence of 0.3 mM Ba^2+^ (*t*_17_=7.317, \*\*\*\**P*<0.0001; unpaired student’s *t* test; n=8-11 recordings/group and N=2-4 male mice/group). **D.** Currents evoked by baclofen (200 μM) in GABA neurons from male GAD67GFP(+) (wild-type), GAD67GFP(+):*Girk1^-/-^*(*Girk1^-/-^*), and GAD67GFP(+):*Girk2^-/-^*(*Girk2^-/-^*) mice. Scale: 25 pA/50 s. **E.** Baclofen-induced currents in GABA neurons from GAD67GFP(+) and GAD67GFP(+):*Girk^-/-^* mice. One-way ANOVA detected a significant difference between groups (F2,23=43.66, *P*<0.0001; n=7-11 recordings/group and N=2-4 male mice/group). *Girk1^-/-^* and *Girk2^-/-^* were significantly different from wild-type (*t*_23_=8.01, \*\*\*\**P*<0.0001; *t*_23_=7.64, \*\*\*\**P*<0.0001). **F.** Baseline rheobase in GABA neurons from GAD67GFP(+) and GAD67GFP(+):*Girk^-/-^* mice. Oneway ANOVA detected no differences between groups (F_2,25_=0.97, *P*=0.39; n=7-11/group and 2-4 male mice/group). **G.** Change in rheobase induced by baclofen (200 μM) in GABA neurons from GAD67GFP(+) and GAD67GFP(+):*Girk^-/-^* mice. One-way ANOVA detected a significant difference between genotypes (F_2,40_=30.29, *P*<0.0001; n=12-16 recordings/group and N=5-6 mice/group). *Girk1^-/-^* and *Girk2^-/-^* were significantly different from wild-type (*t*_40_=5.81, \*\*\*\**P*<0.0001; *t*_40_=7.23, \*\*\*\**P*<0.0001). No main effect of sex was detected (F_1,37_=0.023, *P*=0.88; two-way ANOVA). **H.** Currents evoked by baclofen (200 μM) in GABA neurons from GAD67GFP(+) mice, 1-2 d after repeated saline or cocaine treatment. Currents were reversed by the GABA_B_R antagonist CGP54626 (2 μM). Scale: 25 pA/50 s. **I.** Baclofen-induced currents in GABA neurons from GAD67GFP(+) mice, 1-2 d after repeated saline or cocaine treatment (*t*_32_=0.96, *P*=0.34; unpaired student’s *t* test; n=17 recordings/group and N=5 mice/group). No main effect of sex was detected (F_1,30_=0.0004, *P*=0.98; two-way ANOVA). **J.** RMP in GABA neurons from GAD67GFP(+) mice, 1-2 d after repeated saline or cocaine treatment (*t*_31_=0.54, *P*=0.59; unpaired student’s *t* test; n=16-17 recordings/group and N=5 mice/group). No main effect of sex was detected (F_1,29_=3.4, *P*=0.08; two-way ANOVA). **K.** Rheobase in GABA neurons from GAD67GFP(+) mice, 1-2 d after repeated saline or cocaine treatment (*t*_30_=0.91, *P*=0.37; unpaired student’s *t* test; n=15-17 recordings/group and N=5 mice/group). No main effect of sex was detected (F_1,28_=0.075, *P*=0.79; two-way ANOVA).

### GIRK channel ablation in PL pyramidal neurons

The lack of impact of repeated cocaine on layer 5/6 PL GABA neurons suggests that cocaine exerts a relatively selective impact on adjacent PL pyramidal neurons (40). To probe the behavioral relevance of the GIRK neuroadaptation in layer 5/6 PL pyramidal neurons, we used a neuron-selective viral Cre approach and conditional *Girk1^-/-^*(*Girk1^fl/fl^*) mice (**Fig. 2A**). The CaMKIIa promoter has been used extensively to drive transgene expression in PFC pyramidal neurons (54–57). To evaluate the fidelity of pyramidal neuron targeting with our AAV8/CaMKIIα-based vectors, we infused AAV8-CaMKIIα-mCherry into the PL of GAD67GFP(+) mice. Only a small fraction (4%) of neurons co-expressed GFP and mCherry (**Fig. 2B**), suggesting that AAV8/CaMKIIα-based vectors primarily target pyramidal neurons in the PL.

**Figure 2.**
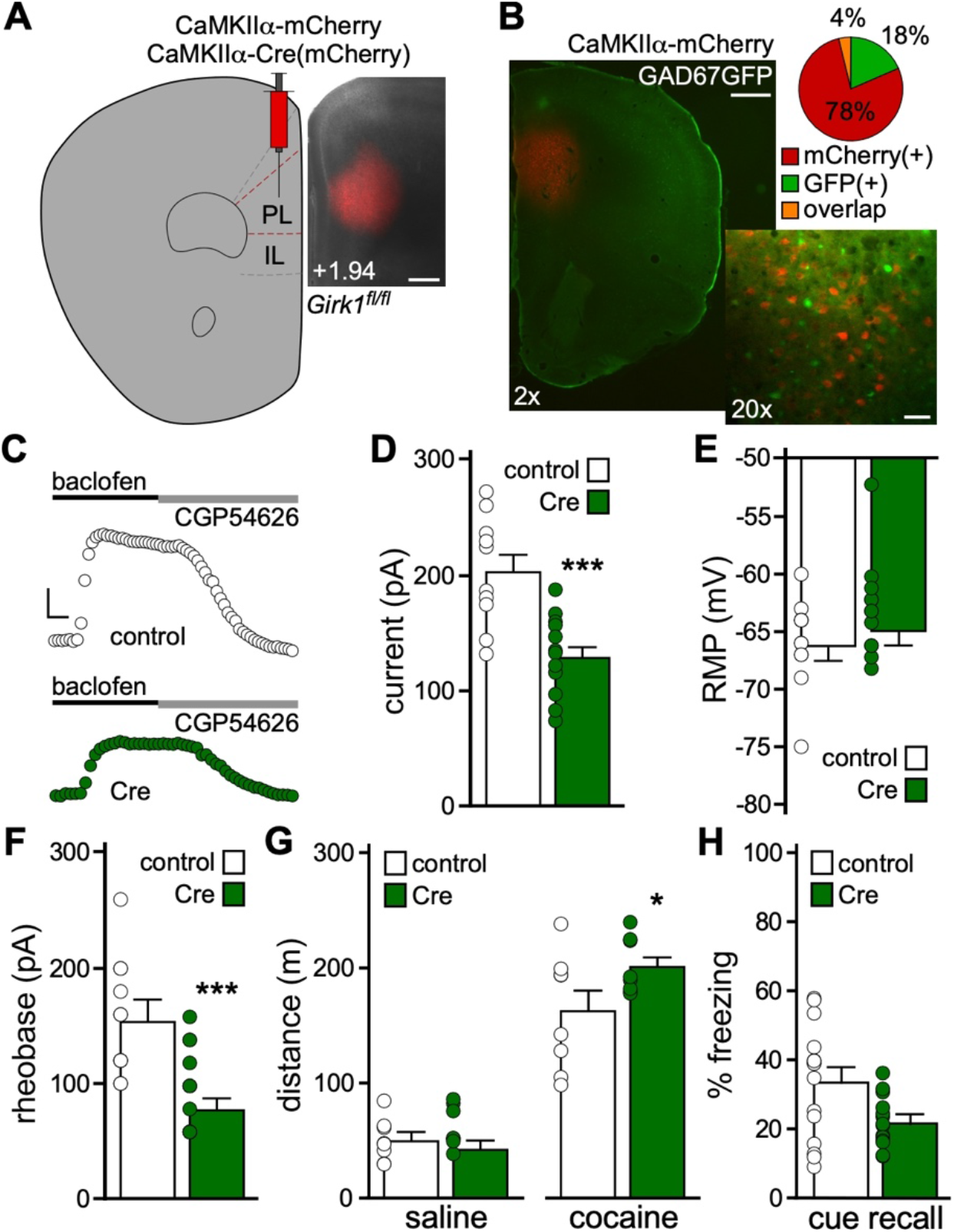
Viral Cre ablation of GIRK channels in PL pyramidal neurons. **A.** Example of viral targeting in a *Girk1^fl/fl^* mouse treated with AAV8-CaMKIIα-Cre(mCherry) vector. Scale: 325 microns. **B.** AAV8-CaMKIIα-mCherry labeling in the PL of a GAD67GFP(+) mouse, and a pie chart depicting percentage of fluorescent neurons expressing mCherry, GFP, or both (overlap) (n=935, 222, 45 neurons, respectively; N=3 mice). Scale: 500 microns (2x)/50 microns (20x). **C.** Currents evoked by baclofen (200 μM) in layer 5/6 PL pyramidal neurons from *Girk1^fl/fl^* mice treated with AAV8-CaMKIIα-Cre(mCherry) or control vector. Currents were reversed by the GABA_B_R antagonist CGP54626 (2 μM). Scale: 50 pA/50 s. **D.** Baclofen-induced currents in layer 5/6 PL pyramidal neurons from *Girk1^fl/fl^* mice treated with AAV8-CaMKIIα-Cre(mCherry) or control vector (*t*20=4.33, \*\*\**P*=0.0003; unpaired student’s *t* test; n=10-12 recordings/group and N=3-6 mice/group). No main effect of sex was detected (F_1,18_=0.15, *P*=0.71; two-way ANOVA). **E.** RMP in layer 5/6 PL pyramidal neurons from *Girk1^fl/fl^* mice treated with CaMKIIα-Cre(mCherry) or control vector (*t*_21_=0.64, *P*=0.53; unpaired student’s *t* test; n=11-12 recordings/group and N=3-6 mice/group). No main effect of sex was detected (F_1,19_=0.079, *P*=0.78; two-way ANOVA). **F.** Rheobase in layer 5/6 PL pyramidal neurons from *Girk1^fl/fl^* mice treated with CaMKIIα-Cre(mCherry) or control vector (*t*_21_=4.32, \*\*\**P*=0.0003; unpaired student’s *t* test; n=11-12/group and N=3-6 mice/group). No main effect of sex was detected (F_1,19_=1.11, *P*=0.31; two-way ANOVA). **G.** Saline- and cocaine-induced motor activity in *Girk1^fl/fl^* mice treated with CaMKIIα-Cre(mCherry) or control vector (N=8 mice/group). A significant interaction between drug and viral vector treatment was detected (F_1,14_=6.30, *P*=0.025; two-way repeated measures ANOVA); pairwise comparison (Bonferroni’s *post hoc* test) revealed a difference between viral treatment groups with respect to cocaine-induced motor activity (*t*_28_=2.51, \**P*=0.036). No main effect of sex (F_1,13_=0.061, *P*=0.81), or sex interactions, were detected (three-way ANOVA). One outlier animal was excluded. **H.** Cue recall test in *Girk1^fl/fl^* mice treated with AAV8-CaMKIIα-Cre(mCherry) or control vector, conducted 2 d after trace fear conditioning (N=14-15 mice/group). No significant difference in freezing was observed during the cue recall test between CaMKIIα-Cre(mCherry)-treated and control mice (U=62.0, *P*=0.063; unpaired non-parametric Mann-Whitney test). No main effect of sex was detected (F_1,25_=1.45, *P*=0.24; two-way ANOVA). One outlier animal was excluded.

AAV8-CaMKIIα-Cre(mCherry) or AAV8-CaMKIIα-mCherry vectors were infused into the PL of *Girk1^fl/fl^* mice. Following a 4-5 wk recovery period, we evaluated the impact of viral Cre and control treatment on mCherry-positive layer 5/6 PL neurons. Viral Cre treatment yielded smaller baclofen-induced currents in these neurons (**Fig. 2C,D**). While loss of GIRK channel activity had no impact on RMP (**Fig. 2E**), rheobase was decreased (**Fig. 2F**), consistent with an increase in excitability.

To assess the behavioral consequences of the manipulation, *Girk1^fl/fl^* mice were infused with CaMKIIa-Cre(mCherry) or control vector, followed by open-field activity assessments. Suppression of GIRK channel activity did not impact distance traveled following saline injection (baseline) but did enhance the motor-stimulatory effect of cocaine (**Fig. 2G**). In a separate cohort, we tested the impact of the manipulation on trace fear conditioning, an associative learning task dependent on PL function (20, 21, 32). Loss of GIRK channel activity in PL pyramidal neurons was associated with decreased cue fear recall, though the difference between Cre-treated and control subjects did not reach statistical significance (*P*=0.063; **Fig. 2H**). Thus, loss of GIRK channel activity in PL pyramidal neurons enhanced the motor-stimulatory effect of cocaine but did not significantly impact baseline activity or trace fear learning.

### Chemogenetic activation of PL pyramidal neurons

As repeated cocaine is associated with multiple adaptations that enhance mPFC pyramidal neuron excitability, we sought to complement the persistent viral Cre manipulation of GIRK channel activity with chemogenetic approaches to acutely enhance PL pyramidal neuron excitability. AAV8-CaMKIIα-hM3Dq(mCherry) or AAV8-CaMKIIα-mCherry vectors were infused into the PL of C57BL/6J mice (**Fig. 3A**). Following a 3-4 wk recovery, we tested whether hM3Dq activation enhanced layer 5/6 PL pyramidal neuron excitability. Bath application of CNO (10 μM) significantly depolarized and decreased the rheobase of hM3Dq(mCherry)-expressing, but not control, PL pyramidal neurons (**Fig. 3B,C**).

**Figure 3.**
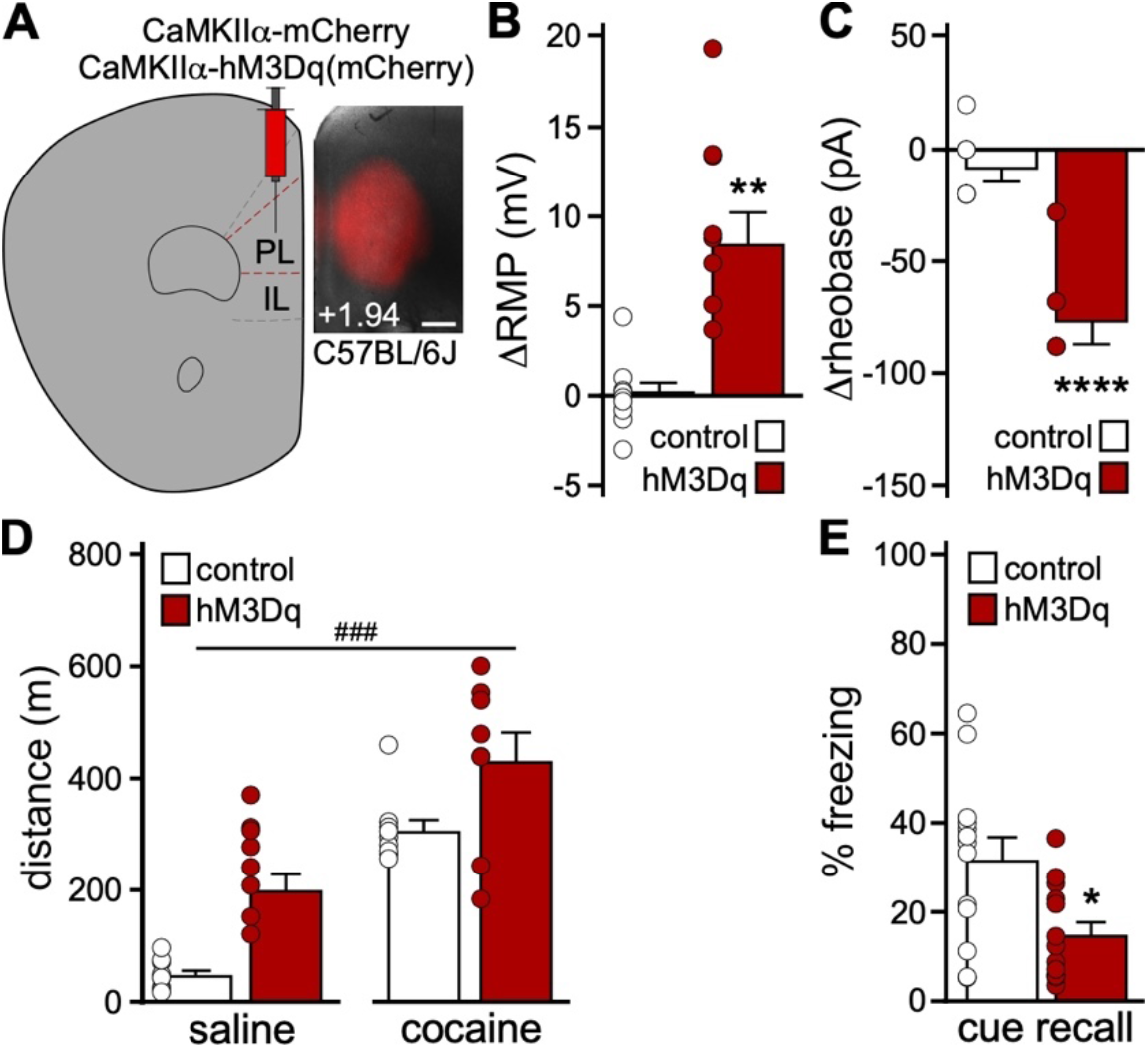
Impact of chemogenetic excitation of PL pyramidal neurons on behavior. **A.** Example of viral targeting in a C57BL/6J mouse treated with AAV8-CaMKIIα-hM3Dq(mCherry). Scale: 325 microns. **B.** Change in RMP induced by CNO (10 μM) in layer 5/6 PL pyramidal neurons from male C57BL/6J mice treated with AAV8-CaMKIIα-hM3Dq(mCherry) or control vector (*t*8.86=4.39, \*\**P*=0.0018; unpaired student’s *t* test with Welch’s correction; n=8-9 recordings/group and N=3 mice/group). **C.** Change in rheobase induced by CNO (10 μM) in layer 5/6 PL pyramidal neurons from male C57BL/6J mice treated with CaMKIIα-hM3Dq(mCherry) or control vector (*t*_12_=5.82, \*\*\*\**P*<0.0001; unpaired student’s *t* test; n=7 recordings/group and N=3 mice/group). **D.** Saline- and cocaine-induced motor activity in male C57BL/6J mice treated with CaMKIIα-hM3Dq(mCherry) or control vector (N=8-9 mice/group), measured 30-min after CNO administration (2 mg/kg IP). Two-way repeated measures ANOVA detected main effects of drug treatment (F_1,15_=68.96, *P*<0.0001) and viral treatment (F_1,15_=19.30, ###*P*=0.0005), but no interaction between drug and viral treatment (F_1,15_=0.17, *P*=0.68). One outlier animal excluded. **E.** Trace fear conditioning in male C57BL/6J mice treated with CaMKIIα-hM3Dq(mCherry) or control vector (N=12-13 mice/group). Lower levels of freezing were observed during cue recall test, conducted 2 d after trace fear conditioning in the presence of CNO (2 mg/kg IP), by CaMKIIα-hM3Dq(mCherry)-treated mice relative to controls (*t*_23_=2.23, \**P*=0.036; unpaired student’s *t* test).

We next examined the impact of chemogenetic excitation of PL pyramidal neurons on motor activity and trace fear conditioning. CNO pre-treatment elevated activity measured after both saline and cocaine injection in hM3Dq(mCherry)-expressing C57BL/6J mice, relative to controls (**Fig. 3D**). Chemogenetic activation of PL pyramidal neurons during trace fear conditioning was associated with lower freezing levels during the subsequent cue recall test (**Fig. 3E**). In a parallel study, we used the well-characterized CaMKIICre line and Cre-dependent AAV vectors to drive expression of hM3Dq(mCherry) or mCherry in PL pyramidal neurons (**Fig. S1A**). In slice validation experiments, CNO (10 μM) depolarized and decreased the rheobase of hM3Dq(mCherry)-expressing layer 5/6 PL neurons (**Fig. S1B,C**). CNO pretreatment elevated motor activity measured after both saline and cocaine injection in hM3Dq(mCherry)-expressing mice (**Fig. S1D**), and chemogenetic excitation of PL pyramidal neurons during trace fear conditioning decreased cue fear recall (**Fig. S1E**). Thus, acute excitation of PL pyramidal neurons increased motor activity at baseline and following cocaine injection, and disrupted trace fear learning.

### Chemogenetic inhibition of PL GABA neurons

Prolonged cocaine exposure reduces GABAergic neurotransmission in PL pyramidal neurons (40, 43, 44), which should indirectly enhance PL pyramidal neuron excitability. Indeed, chemogenetic inhibition of layer 5/6 PL GABA neurons decreased the frequency of spontaneous inhibitory postsynaptic currents (sIPSCs) in adjacent pyramidal neurons (**Fig. S2**). To mimic reduced GABAergic input to PL pyramidal neurons in drug-naïve C57BL/6J mice, we used a viral chemogenetic approach involving the forebrain GABAergic neuron promoter/enhancer mDlx (58) to acutely inhibit PL GABA neurons (**Fig. 4A**). To test whether AAV8/mDlx-based vectors selectively targeted PL GABA neurons, we infused AAV8-mDlx-mCherry into the PL of GAD67GFP(+) mice. A large majority (76%) of PL neurons co-expressed GFP and mCherry, and a small fraction (7%) expressed only mCherry (**Fig. 4B**). Thus, AAV8/mDlx-based vectors afford relatively selective access to mouse PL GABA neurons. Notably, CNO hyperpolarized and increased the rheobase of hM4Di(mCherry)-expressing layer 5/6 PL GABA neurons in C57BL/6J mice (**Fig. 4C,D**).

**Figure 4.**
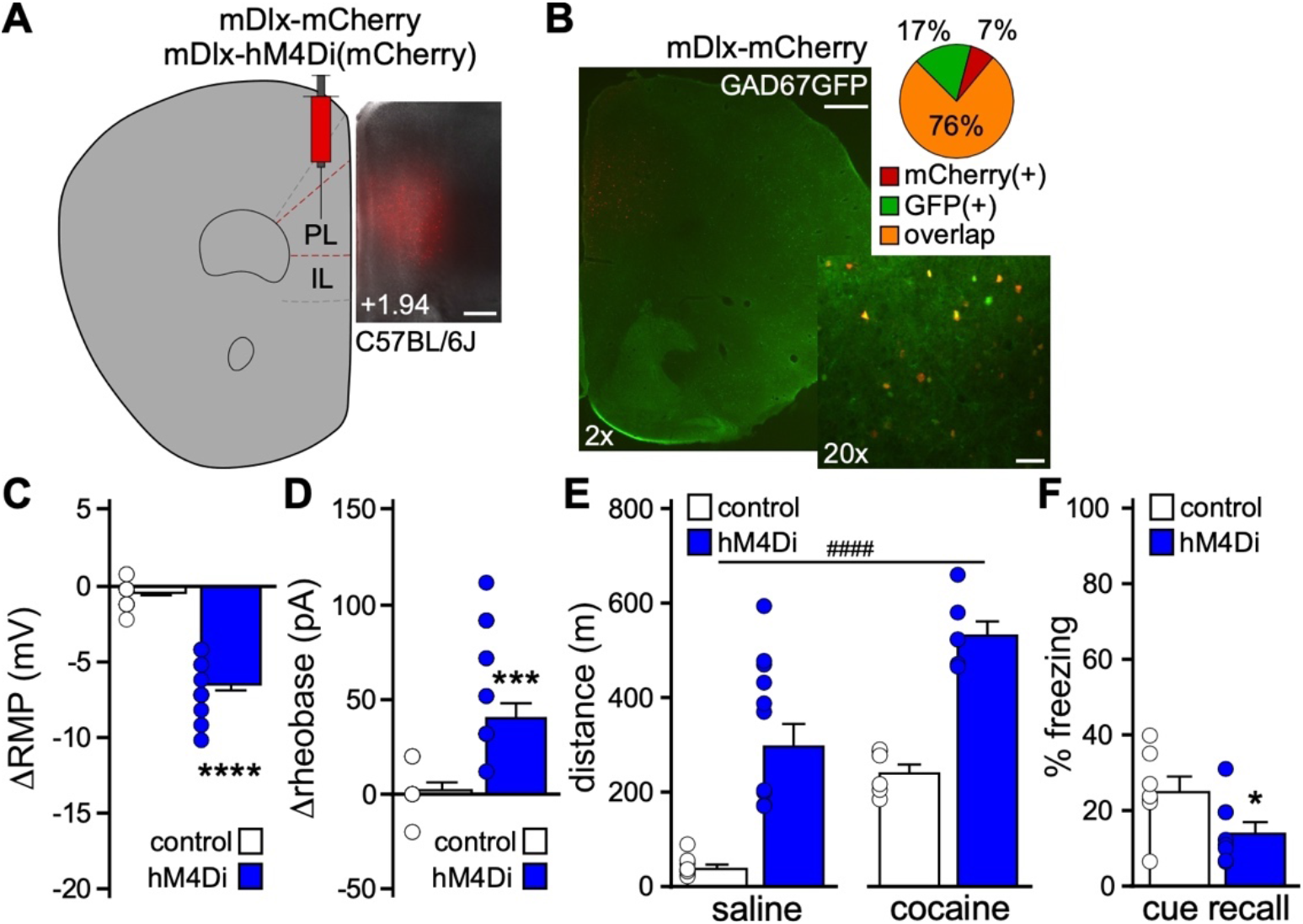
Impact of chemogenetic inhibition of PL GABA neurons on behavior. **A.** Example of viral targeting in a C57BL/6J mouse treated with AAV8-mDlx-mCherry. Scale: 325 microns. **B.** AAV8-mDlx-mCherry labeling in the PL of GAD67GFP(+) mouse, and pie chart depicting percent of fluorescent neurons expressing mCherry, GFP, or both (overlap) (n=26, 62, 286 neurons, respectively; N=3 mice). Scale bars: 500 microns (2x)/50 microns (20x). **C.** Change in RMP induced by CNO (10 μM) in layer 5/6 PL GABA neurons from male C57BL/6J mice treated with AAV8-mDlx-hM4Di(mCherry) or control vector (*t*_18.07_=11.08, \*\*\*\**P*<0.0001; unpaired student’s *t* test with Welch’s correction; n=14 recordings/group and N=6 mice/group). **D.** Change in rheobase induced by CNO (10 μM) in layer 5/6 PL GABA neurons from male C57BL/6J mice treated with AAV8-mDlx-hM4Di(mCherry) or control vector (*t*_21.45_=4.374, \*\*\**P*=0.0003; unpaired student’s *t* test with Welch’s correction; n=13-16 recordings/group and N=6 mice/group). One outlier data point excluded. **E.** Saline- and cocaine-induced motor activity in separate cohorts of male C57BL/6J mice treated with mDlx-hM4Di(mCherry) or control vector (N=6-10 mice/group), measured 30-min after systemic CNO administration (2 mg/kg IP). Two-way ANOVA detected main effects of drug (F_1,27_=38.64, *P*<0.0001) and viral treatment (F_1,27_=61.19, ####*P*<0.0001), but no interaction between drug and viral treatment (F_1,27_=0.25, *P*=0.62). One outlier animal was excluded. **F.** Trace fear conditioning in male C57BL/6J mice treated with mDlx-hM4Di(mCherry) or control vector (N=7-8 mice/group). Lower levels of freezing were observed during the cue recall test, conducted 2 d after trace fear conditioning in the presence of CNO (2 mg/kg IP), by mDlx-hM4Di(mCherry)-treated mice relative to controls (*t*_13_=2.20, \**P*=0.047; unpaired student’s *t* test).

We next examined the impact of chemogenetic inhibition of PL GABA neurons on motor activity and trace fear conditioning in C57BL/6J mice. CNO pre-treatment elevated motor activity measured after both saline and cocaine injection in hM4Di(mCherry)-treated subjects, compared to controls (**Fig. 4E**), and chemogenetic inhibition of PL GABA neurons during trace fear conditioning was associated with decreased cue fear recall (**Fig. 4F**). Thus, chemogenetic inhibition of PL GABA neurons, like chemogenetic excitation of PL pyramidal neurons, increased motor activity at baseline and following cocaine injection, and disrupted trace fear learning.

### Chemogenetic activation of distinct PL projections

We next used a projection-specific viral chemogenetic approach to manipulate PL neurons projecting to the BLA, NAc, or VTA (59–62). These brain regions were selected because they receive glutamatergic input from the PL and given their involvement in fear learning and/or motor activity (5, 63–68). We infused an AAV2retro-based (69) Cre vector (AAV2retro-hSyn-Cre-GFP) into the downstream target of interest, and a Cre-dependent vector (AAV8-hSyn-DIO-hM3Dq(mCherry) or AAV8-hSyn-DIO-mCherry) into the PL (**Fig. 5A,B,E,H**). The impact of chemogenetic excitation of each PL projection was first assessed using trace fear conditioning, 4-5 wk after surgery. While excitation of BLA- or NAc-projecting PL pyramidal neurons during trace fear conditioning did not affect cue fear recall (**Fig. 5C,F**), excitation of VTA-projecting PL pyramidal neurons disrupted cue fear learning (**Fig. 5I**).

**Figure 5.**
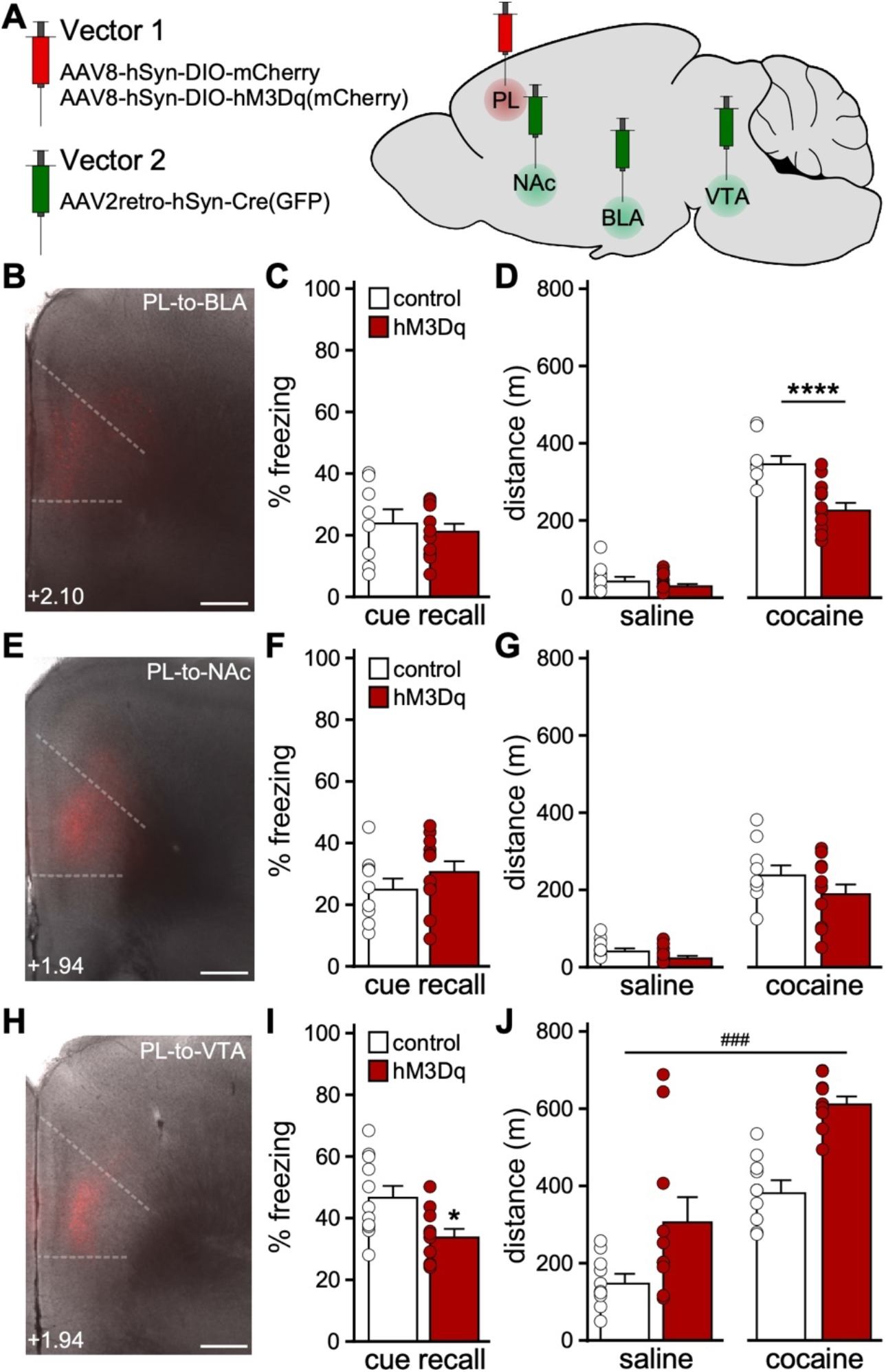
Impact of chemogenetic excitation of distinct PL projections on behavior. **A.** Projection-specific chemogenetic approach involving a Cre-dependent Vector 1 (AAV8-hSyn-DIO-hM3Dq(mCherry) or AAV8-hSyn-DIO-mCherry) infused into the PL, and AAV2retro-hSyn-Cre(GFP) was infused into the BLA, NAc, or VTA. **B.** Cre-dependent mCherry expression in the PL of a C57BL/6J mouse treated with intra-PL AAV8-hSyn-DIO-mCherry and intra-BLA AAV2retro-hSyn-Cre(GFP). Scale: 325 microns. **C.** Trace fear conditioning in male C57BL/6J mice treated with intra-PL DIO-hM3Dq(mCherry) or control vector, and intra-BLA AAV2retro-hSyn-Cre(GFP) (N=8-11 mice/group). No difference in freezing was observed during the cue recall test, conducted 2 d after trace fear conditioning in the presence of CNO (2 mg/kg IP), by DIO-hM3Dq(mCherry)-treated mice relative to controls (*t*_17_=0.543, *P*=0.59; unpaired student’s *t* test). **D.** Saline- and cocaine-induced motor activity in male C57BL/6J mice treated with intra-PL DIO-hM3Dq(mCherry) or control vector, and intra-BLA AAV2retro-hSyn-Cre(GFP) (N=8-11 mice/group), measured 30-min after CNO administration (2 mg/kg IP). A significant interaction between drug and viral treatment was detected (F_1,17_=13.17, *P*=0.0021; two-way repeated measures ANOVA); pairwise comparison (Bonferroni’s *post hoc* test) revealed a difference between viral treatment groups with respect to cocaine-induced motor activity (*t*_34_=5.314, \*\*\*\**P* <0.0001). **E.** Cre-dependent mCherry expression in the PL of a C57BL/6J mouse treated with intra-PL AAV8-hSyn-DIO-mCherry and intra-NAc AAV2retro-hSyn-Cre(GFP). Scale: 325 microns. **F.** Trace fear conditioning in male C57BL/6J mice treated with intra-PL DIO-hM3Dq(mCherry) or control vector, and intra-NAc AAV2retro-hSyn-Cre(GFP) (N=9-11 mice/group). No difference in freezing was observed during the cue recall test, conducted 2 d after trace fear conditioning in the presence of CNO (2 mg/kg IP), by DIO-hM3Dq(mCherry)-treated mice relative to controls (*t*_18_=1.109, *P*=0.28; unpaired student’s *t* test). **G.** Saline- and cocaine-induced motor activity in male C57BL/6J mice treated with intra-PL DIO-hM3Dq(mCherry) or control vector, and intra-NAc AAV2retro-hSyn-Cre(GFP) (N=9-11 mice/group), measured 30-min after CNO administration (2 mg/kg IP). Two-way repeated measures ANOVA detected a main effect of drug treatment (F_1,18_=99.73, *P*<0.0001), but no main effect of viral treatment (F_1,18_=2.795, *P*=0.11), and no interaction between drug and viral treatment (F_1,18_=0.74, *P*=0.40). **H.** Cre-dependent mCherry expression in the PL of a C57BL/6J mouse treated with intra-PL AAV8-hSyn-DIO-mCherry and intra-VTA AAV2retro-hSyn-Cre(GFP). Scale: 325 microns. **I.** Trace fear conditioning in male C57BL/6J mice treated with DIθ-hM3Dq(mCherry) or control vector, and intra-VTA AAV2retro-hSyn-Cre(GFP) (N=10-11 mice/group). DIO-hM3Dq(mCherry)-treated mice exhibited reduced freezing relative to controls during the cue recall test (*t*_19_=2.667, \**P*=0.0152; unpaired student’s *t* test), conducted 2 d after trace fear conditioning in the presence of CNO (2 mg/kg IP). **J.** Saline- and cocaine-induced motor activity in male C57BL/6J mice treated with intra-PL DIO-hM3Dq(mCherry) or control vector, and intra-VTA AAV2retro-hSyn-Cre(GFP) (N=10 mice/group), measured 30-min after systemic CNO administration (2 mg/kg IP). Two-way repeated measures ANOVA detected main effects of drug treatment (F_1,18_=63.05, *P*<0.0001) and viral treatment (F_1,18_=20.77, ###*P*=0.0002), but no interaction between drug and viral treatment (F_1,18_=1.012, *P*=0.33). One outlier animal excluded.

We next assessed the impact of exciting each PL projection on motor activity. Chemogenetic excitation of BLA-projecting PL pyramidal neurons did not impact saline-induced activity (**Fig. 5D**, left), but did suppress cocaine-induced activity (**Fig. 5D**, right). Chemogenetic excitation of NAc-projecting PL pyramidal neurons had no effect on saline- or cocaine-induced motor activity (**Fig. 5G**). Excitation of VTA-projecting PL pyramidal neurons enhanced activity measured after saline or cocaine injection (**Fig. 5J**). Thus, acute excitation of VTA-projecting PL pyramidal neurons recapitulated the motor activity and trace fear learning phenotypes seen with comprehensive excitation of PL pyramidal neurons.

## DISCUSSION

Previously, we reported that repeated cocaine exposure increased layer 5/6 PL pyramidal neuron excitability, likely due to a suppression of GIRK-dependent signaling (40). These adaptations were evident even during early withdrawal (1-2 d after the last cocaine injection) and were not seen in adjacent layer 2/3 pyramidal neurons or layer 5/6 IL pyramidal neurons. Using an identical cocaine treatment regimen and recording timeline, we found that repeated cocaine does not impact GIRK-dependent signaling in, or excitability of, layer 5/6 PL GABA neurons. As some cocaine-induced adaptations are evident at earlier (70) or later (71) withdrawal timepoints, our cocaine treatment regimen may evoke adaptations in layer 5/6 PL GABA neurons outside the 1-2 d withdrawal window. It is also possible that repeated cocaine provokes adaptations in distinct interneuron sub-populations (70).

RNAi-based suppression of GIRK channel expression in the PL enhanced the motor-stimulatory effect of cocaine (40). Here, we show that selective suppression of GIRK channel activity in PL pyramidal neurons recapitulates this phenotype. This finding aligns with other reports implicating the PL and GABA_B_R-dependent signaling in locomotor sensitization. For example, PL lesions blocked the induction and expression of cocaine-induced locomotor sensitization (17–19), and baclofen infusion into the mPFC blocked acute cocaine-induced locomotion and induction of locomotor sensitization without affecting basal activity (72, 73). Collectively, these lines of evidence suggest that the cocaine-induced suppression of GIRK-dependent signaling in PL pyramidal neurons contributes to locomotor sensitization.

The mPFC regulates cognitive functions (20, 21), including trace fear learning (28, 31, 32). Persistent firing in the PL during the trace interval is critical for acquisition of trace fear learning (23–32), and optogenetic silencing of the PL during the trace interval precludes fear learning (32). Our chemogenetic data show that acute excitation of PL pyramidal neurons during conditioning can also disrupt fear learning. Disruptions in trace fear learning during conditioning may reflect impairments in attention and/or working memory (20), processes that are altered by psychostimulant exposure in humans (74–76), non-human primates (77), and rodents (78, 79).

Prolonged exposure to cocaine persistently elevates PL pyramidal neuron excitability (35, 40–42, 80) and reduces GABAergic neurotransmission in these neurons (40, 43, 44). In an attempt to mimic these adaptations in drug-naïve mice, we used three distinct approaches: viral Cre ablation of GIRK channels, chemogenetic activation of PL pyramidal neurons, and chemogenetic inhibition of PL GABA neurons. While viral Cre ablation of GIRK channel activity in PL pyramidal neurons did not impact saline-induced motor activity, direct or indirect chemogenetic excitation of PL pyramidal neurons did. The outcomes were surprising given that hM3Dq activation in PL pyramidal neurons did not alter open field activity (54, 56). These differences may relate to the scope of viral targeting, open field test duration, and/or prior chemogenetic activation and behavioral testing history of subjects in the different studies. In line with our results, disinhibiting PL neurons by antagonizing GABA_A_Rs or blocking GABA synthesis increased locomotion in 30-min open field tests (81, 82).

hM3Dq activation in PL pyramidal neurons enhanced key measures of neuronal excitability (rheobase and RMP), enhanced motor activity measured after saline or cocaine injection, and disrupted trace fear learning. In contrast, GIRK ablation increased PL pyramidal neuron excitability (rheobase but not RMP), enhanced cocaine-induced but not baseline activity, and evoked a non-significant decrease in trace fear learning. Why do persistent (GIRK ablation) and acute (chemogenetic) manipulations targeting PL pyramidal neurons yield overlapping but distinct behavioral outcomes? We speculate that ablation of GIRK channels, predominantly located in the somatodendritic compartment (83, 84), preferentially impact somatodendritic physiology of PL pyramidal neurons, whereas hM3Dq activation exerts a multi-faceted influence on intracellular signaling in somatodendritic/postsynaptic and axonal/presynaptic compartments (85). Moreover, the persistent suppression of GIRK channel activity may promote compensatory adaptations not seen in acute chemogenetic models.

For our projection-specific manipulations, we targeted brain regions that receive glutamatergic input from the PL and regulate fear learning and/or motor activity. The NAc has been implicated in fear learning (65) and the acute motor-stimulatory effect of cocaine (86), and mPFC inputs are involved in the development and expression of locomotor sensitization (5). Nevertheless, we did not see any impact of exciting NAc-projecting PL pyramidal neurons on fear learning, or motor activity. Consistent with the latter finding, optogenetic stimulation of dorsal mPFC-to-NAc projections did not alter movement velocity in mice (62). Similarly, despite evidence that activity in the PL and BLA are necessary for trace fear learning (32, 66, 68), we found that exciting BLA-projecting PL pyramidal neurons was without effect. This result is consistent with a study showing that optogenetic excitation of dorsal mPFC-amygdala projections did not affect acquisition of cue fear in a delay fear conditioning model (87). Interestingly, while exciting BLA-projecting PL pyramidal neurons did not affect basal locomotion, in line with similar reports (87, 88), this manipulation suppressed cocaine-induced activity. Given data showing that reversible inactivation (muscimol/baclofen) of the amygdala enhanced cocaine-induced locomotion (63), our findings suggest that BLA neuron excitability is negatively correlated with cocaine-induced motor activity and subject to modulation via PL glutamate.

The VTA plays a significant role in motor activity and locomotor sensitization (5, 67). Chemogenetic inhibition of VTA DA neurons reduced basal and cocaine-induced locomotion (89), while exciting VTA DA neurons, or the VTA-to-NAc projection, elevated basal locomotion (67, 89–92). mPFC pyramidal neurons synapse onto VTA DA neurons (60, 93, 94) and activation of the mPFC (95, 96) or intra-VTA infusion of glutamatergic agonists (97, 98) induces burst-spiking of VTA DA neurons *in vivo*. mPFC stimulation also enhances DA release in the NAc (95, 99, 100), suggesting that exciting the PL projection to the VTA directly activates the mesolimbic DA pathway. Indeed, an excitatory monosynaptic projection from the mPFC to NAc-projecting VTA DA neurons has been reported (101). VTA-projecting mPFC neurons exhibit cocaine-induced plasticity that facilitates glutamate release in the VTA and contributes to addictive behaviors, including behavioral sensitization (5, 40). We report here that acute stimulation of PL projections to the VTA enhances motor activity measured following saline or cocaine injection. The VTA also regulates aversive learning (64), and optogenetic inhibition of VTA DA neurons during footshock (but not auditory cue) presentation enhanced cue fear recall (102). We show that acute excitation of VTA-projecting PL pyramidal neurons during trace fear conditioning disrupts cue fear recall. As the PL-to-VTA projection is subject to cocaine-induced plasticity (40), it is tempting to speculate that some of the dysregulation of learning/memory processes seen following chronic cocaine exposure is linked in part to enhanced excitability of VTA-projecting PL pyramidal neurons (22).

In summary, we show that distinct manipulations of PL pyramidal neuron excitability in drug-naïve mice exert overlapping but distinct consequences on behaviors relevant to addiction. Our work further suggests that enhanced excitability of the glutamatergic PL projection to the VTA pre-sensitizes mice to the motor-stimulatory effect of cocaine and disrupts associative fear learning. As such, interventions that suppress the excitability of this microcircuit may prove useful for suppressing problematic behaviors linked to chronic cocaine intake.

## ACKNOWLEDGEMENTS

The authors would like to thank Zhilian Xia, Nicholas Carlblom, Hannah Oberle, and Mehrsa Zahiremami for exceptional care of the mouse colony. This work was supported by NIH grants to KW (MH061933, DA034696, AA027544), MET (AA025978), and TRR (DA007234), as well as a Wallin Neuroscience Discovery Fund Award (KW) and a Doctoral Dissertation Fellowship (BNV) from the University of Minnesota. The authors declare no competing interests.

## Notes

### Competing Interest Statement

The authors have declared no competing interest.

