## Supplemental Information for "Divergent behavioral consequences of manipulations enhancing pyramidal neuron excitability in the prelimbic cortex"

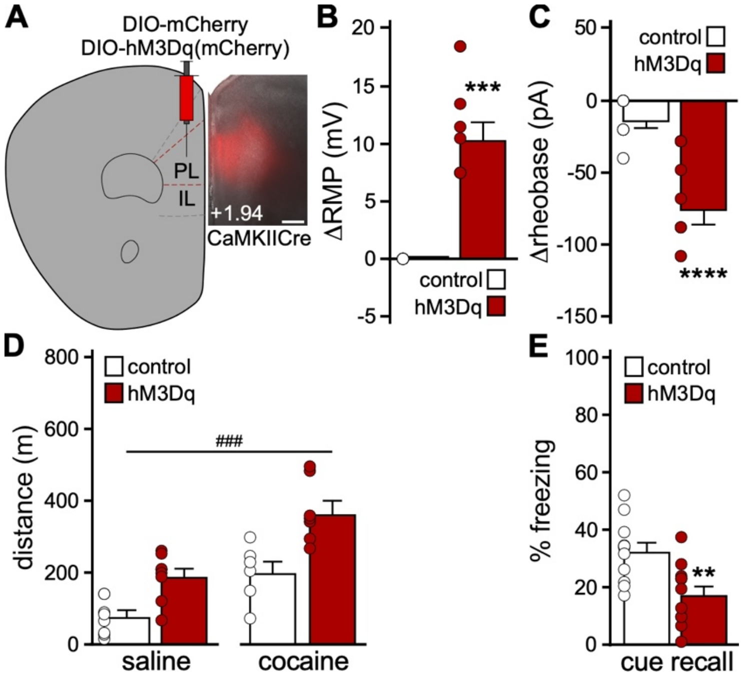


**Figure S1. Impact of chemogenetic excitation of PL pyramidal neurons on behavior**

**A.** Example of viral targeting in a CaMKIICre(+) mouse treated with AAV8-hSyn-DIO-mCherry. Scale: 325 microns.

**B.** Change in RMP induced by CNO (10 μM) in layer 5/6 PL pyramidal neurons from CaMKIICre(+) mice treated with AAV8-hSyn-DIO-hM3Dq(mCherry) or control vector ($t_7$=6.34, ****P*=0.0004; unpaired student's *t* test with Welch's correction; n=8-11 recordings/group and N=3-5 mice/group). No main effect of sex was detected ($F_{1,15}$=1.94, *P*=0.18; two-way ANOVA).

**C.** Change in rheobase induced by CNO (10 μM) in layer 5/6 PL pyramidal neurons from CaMKIICre(+) mice treated with DIO-hM3Dq(mCherry) or control vector ($t_{14.26}$=5.58, *****P*<0.0001; unpaired student's *t* test with Welch's correction; n=11 recordings/group and N=3-5 mice/group). No main effect of sex was detected ($F_{1,18}$=0.82, *P*=0.38; two-way ANOVA).

**D.** Saline- and cocaine-induced motor activity in CaMKIICre(+) mice treated with DIO-hM3Dq(mCherry) or control vector (N=6-8 mice/group), measured 30-min after systemic CNO administration (2 mg/kg IP). Mixed effects analysis revealed main effects of drug treatment ($F_{1,11}$=66.58, *P*<0.0001) and viral treatment ($F_{1,14}$=19.64, ###*P*=0.0006), but no interaction between drug and viral treatment ($F_{1,11}$=1.83, *P*=0.20). No main effect of sex ($F_{1,12}$=1.12, *P*=0.31), or sex interactions, were detected (mixed-effects model). One outlier animal was excluded.

**E.** Trace fear conditioning in CaMKIICre(+) (hM3Dq) and CaMKIICre(-) (control) mice treated with DIO-hM3Dq(mCherry) vector (N=11 mice/group). Lower levels of freezing were observed during the cue recall test ($t_{20}$=3.23, ***P*=0.0042; unpaired student's *t* test), conducted 2 after trace fear conditioning in the presence of CNO (2 mg/kg IP), by CaMKIICre(+) mice relative to CaMKIICre(-) controls. No main effect of sex was detected ($F_{1,18}$=1.48, *P*=0.24; two-way ANOVA).

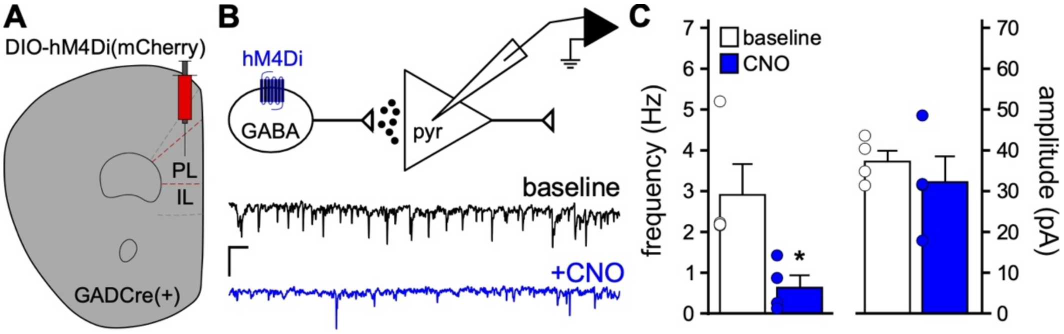


**Figure S2. Chemogenetic inhibition of PL GABA neurons disinhibits adjacent pyramidal neurons**

**A.** GADCre(+) mice were treated with intra-PL AAV8-hSyn-DIO-hM4Di(mCherry) vector.

**B.** Spontaneous inhibitory postsynaptic currents (sIPSCs) were recorded in layer 5/6 PL pyramidal neurons from GADCre(+) mice treated with DIO-hM4Di(mCherry) vector ($V_{hold}$=-70 mV), before (baseline) and after bath application of CNO (10 $\mu$M). Scale = 20 pA/1 s.

**C.** sIPSC frequency and amplitude in layer 5/6 PL pyramidal neurons, measured before and after CNO (10 $\mu$M) application in slices from GADCre(+) mice treated with DIO-hM4Di(mCherry) vector. CNO reduced sIPSC frequency (left: $t_3$=4.32, *$P$=0.023; paired student's $t$ test) but not amplitude (right: $t_3$=0.90, $P$=0.43; paired student's $t$ test); n=4 recordings/group and N=2 male mice/group.
